# The geography of nocturnality – emergence patterns of bats on a latitudinal gradient

**DOI:** 10.64898/2026.08.26.747239

**Authors:** Samanvitha Santusht, Mari Aas Fjelldal, Rohit Chakravarty, Giulliana Appel, Paulo Bobrowiec, Thomas M. Lilley

## Abstract

Animals often attempt to resolve the trade-off between foraging and predation by regulating their activity timings and patterns. Driven by individual energetic requirements and the local environment, onset of activity in a species varies across space and time. These differences are particularly enhanced on a latitudinal gradient, influenced by varying lengths of daylight, temperature and seasonality. For nocturnal mammals like bats, such patterns translate to shorter activity windows at higher latitudes, both on a nightly and a seasonal basis. This constructs the timing of activity onset as a crucial variable for a successful night of foraging. While roost exit timings and influence of local conditions on bat activity have been extensively studied across species inhabiting a variety of locations around the world, our understanding of bats’ emergence behaviour on a global scale remains limited. To investigate emergence patterns in bats across a latitudinal gradient, we conducted a meta-analysis spanning 92 species across 13 Chiropteran families. We evaluated how timing and luminosity (i.e., sun altitude) at exit varied across latitudes and species. Results revealed that bats across the globe emerged under conserved thresholds of brightness, a pattern that was facilitated by a temporal delay in roost exit at higher latitudes. By maintaining emergences within the confines of nautical twilight (sun altitude between 0° to -12°), bats in known predatory environments appear to balance the dilemma between risk and food. This pattern in brightness at emergence was mirrored across different social structures (maternal v. non-maternal colonies) and two broad dietary types (frugivores and insectivores). Furthermore, our models indicated that while local conditions dictate final exit decisions, shared ancestry was potentially influencing the extent of plasticity seen in emergence patterns. Lastly, on comparing exit patterns of bats in regions with and without predators, we found that species in locations without any known predators emerged from their roosts before sunset, and under consistently brighter conditions.

**Graphical abstract:** 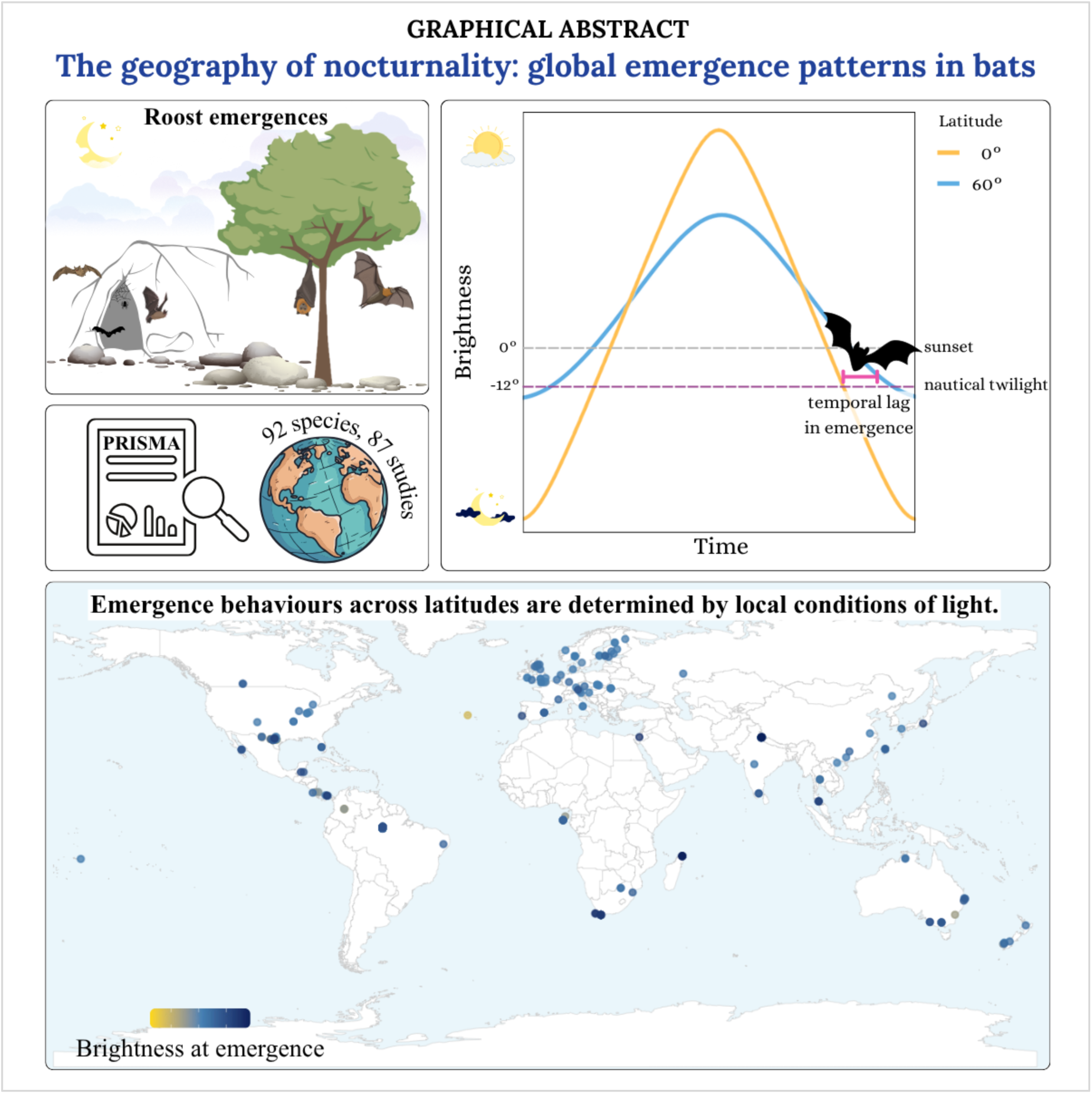

## Introduction

The costs and benefits of activity vary temporally, which is reflected in the distribution of activity in animals across the diel cycle (Bradshaw & Holzapfel, 2007). The drivers of diel activity patterns include resource availability (e.g. food, water, shelter), habitat quality, predation risks, thermoregulatory constraints, mating or socializing opportunities, and inter- or intra-specific competition (Hut et al., 2012; Lima & Dill, 1990; Vallejo-Vargas et al., 2022). Species can be classified into four main groups based on the temporal concentration of their activity; diurnal (predominantly active at day), nocturnal (active at night), crepuscular (active at twilight), and cathemeral (no pronounced activity peaks based on photoperiod) (Aschoff, 1989; Refinetti, 2008). Although diel activity patterns can vary at the individual level within these groups (Hertel et al., 2017; Refinetti, 2008), species with strong diel drivers (e.g., high predation risk) might express more similarities in their activity patterns.

One such taxon is Chiroptera; forming the second most species-rich mammalian order, with more than 1500 species globally (Simmons & Cirranella, 2026), nearly all bats are exclusively nocturnal. Explanations for this taxon-wide adaptation include three main theories that focus on light-dependent costs of activity: predation risk by avian predators, temporal competition for resources by diurnal birds, and risk of hyperthermia (Speakman, 1995). Although no strong support exists yet, many studies point towards predation risk as an important driver (Arndt et al., 2018; Lima & O’Keefe, 2013; Rydell & Speakman, 1995; Speakman, 1991), a hypothesis that is strengthened by greater observation-frequencies of day-flying bats in predator-free environments (Chua & Aziz, 2019; Leonardo & Medeiros, 2011; Rydell & Speakman, 1995). Regardless of the cause, bats worldwide generally express aversion against flying in daylight and spend the daytime in roosts (Kunz, 1982), only emerging from these sometime within their active window, i.e., between sunset and sunrise (Erkert, 1982).

During this timeframe, bats forage, drink, swarm and investigate new roosts and/or hibernation sites (Altringham, 1996; Fraser & McGuire, 2023). To maximise the time available for activity, bats can either advance their dusk emergence or delay their dawn returns. Globally, and across dietary niches, bats are found to be most active in the first few hours following sunset, attempting to overlap their activity with periods of maximum food availability (Acharya et al., 2015; Cano & Murillo-García, 2021; Lilley, Fjelldal, et al., 2026). Therefore, pushing exit times closer toward sunset could allow for greater foraging benefits. However, under such circumstances, bats would be active under brighter conditions, a period marked by higher probabilities of diurnal predator activity. Thus individuals need to balance the trade-off between exiting early and risking predation or emerging under darker conditions with reduced foraging time. This compromise is particularly pronounced in lactating females, who have higher energy demands and exit earlier than their counterparts (Fjelldal et al., 2025; Lilley, Fjelldal, et al., 2026). Based on the influence emergence patterns have on immediate fitness of individuals, this behaviour has been studied extensively.

A commonly reported emergence parameter to measure these trade-offs is timing emergence in relation to sunset, i.e., how many minutes before or after sunset did activity began. Ultimately, time since sunset acts as a proxy for the conditions of brightness at emergence; however, studies directly quantifying light conditions at emergence are few (but see Feng et al., 2022; Fjelldal et al., 2025; Welbergen, 2008), and difficult to compare due to light-measurements being easily influenced by local factors such as vegetation cover, topography or the angle/direction/type of light-logger used. Consequently, time since sunset has been the common measure for evaluating and comparing emergence behaviour in bats. The actual effect of light on timing of emergence on a global, taxon-wide scale therefore remains unresolved.

An important point to consider with brightness thresholds on a global scale is the rate of change of light conditions, both between and within days, across latitudes and seasons. While photoperiod remains highly consistent at the equator, it shows considerable seasonal variation with increasing latitude, where night-length decreases and duration of twilight increases as we approach summer solstice (fig 1a). This pattern is viewed as a direct consequence of the axial tilt of Earth, with the Sun dipping just below the horizon during summer season at the higher latitudes (fig 1b).

**Fig. 1:**
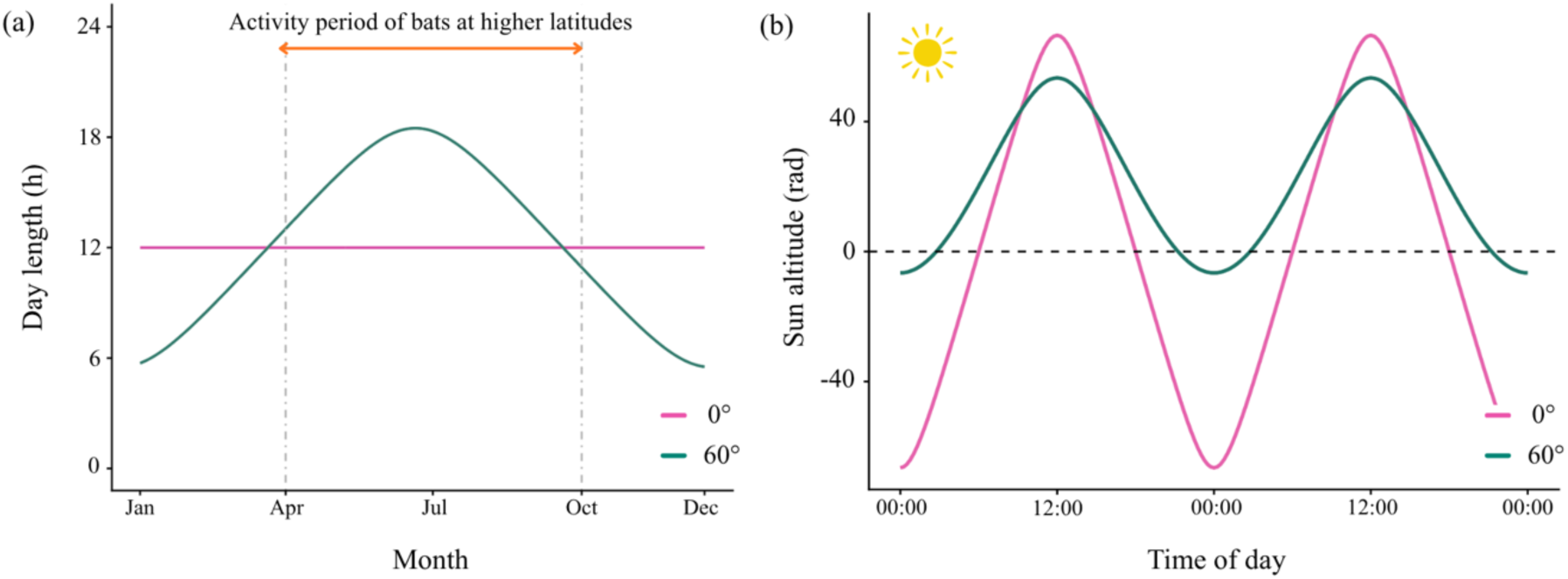
(a) Annual variation in day length for 0° and 60° latitudes. (b) Solar altitude during summer solstice, at 0° and 60° latitudes. Dotted horizontal line represents the horizon (sunset / sunrise).

For bats at these latitudes, the short nights and extended twilight (fig. 1b) should result in them emerging under brighter conditions than bats closer to the equator. Regardless, irrespective of geography, bats across the globe continue to maintain their nocturnal nature, (L. Feng et al., 2022; Frafjord, 2021; Rydell, 1992). While variation in local environmental conditions (e.g. temperature, precipitation, wind speed) are known to impact emergence times (Duvergé et al., 2000; Geipel et al., 2019; Lilley, Vesterinen, et al., 2026a; Russo et al., 2007), we lack knowledge on the influence of one of the most important abiotic environmental drivers, i.e., luminosity. Does tolerance toward brighter conditions increase as we move to higher latitudes, or do bats share a common, global light-threshold?

To understand how latitudinal variations in twilight influence emergence behaviours of bats across different diets and reproductive phases, we conducted a meta-analysis. We aimed to investigate roost-emergence patterns on (1) a temporal scale (i.e., time since sunset at exit), and (2) a luminance range (i.e., corresponding sun altitude at exit). By modelling time and conditions of light against a latitudinal gradient, we focused on testing if species’ geographies influence their light tolerance. Accounting for the foraging pressures and prolonged durations of twilight at higher latitudes, we hypothesized that temperate species would show temporally later emergences than their tropical counterparts. However, given the lighter environment, we expected this emergence to occur under relatively brighter conditions than that of equatorial colonies.

## Methods

### 2.1 Literature search

To compile data on emergence time across bat species, we carried out a literature review, conducted in accordance with the protocols set in PRISMA (Preferred Reporting Items for Systematic Reviews and Meta-Analyses, (Page et al., 2021)). We looked for publications in the Web of Science and Google Scholar’s advanced search databases, followed by specific searches in 3 data repositories – Dryad, Figshare and Zenodo. The following string was used in the searches: “bat*” OR “chiroptera*” AND “emergence” OR “activity pattern*” OR “roost exit*” OR “forag*” AND “roost”. The final search was carried out in January 2026.

Articles were screened based on relevant titles and abstracts. Documents that did not mention or include emergence times or roost exiting behaviour in bats were excluded. Studies that met these basic criteria were read, assessed and retained based on two primary conditions: (1) direct report of mean emergence times and (2) availability of raw emergence time data. Additionally, studies that lacked geographical coordinates and/or temporal data were also filtered out. The final tally of publications included in this meta-analysis came to 87 studies (fig. 2), spanning 101 species.

**Fig. 2:**
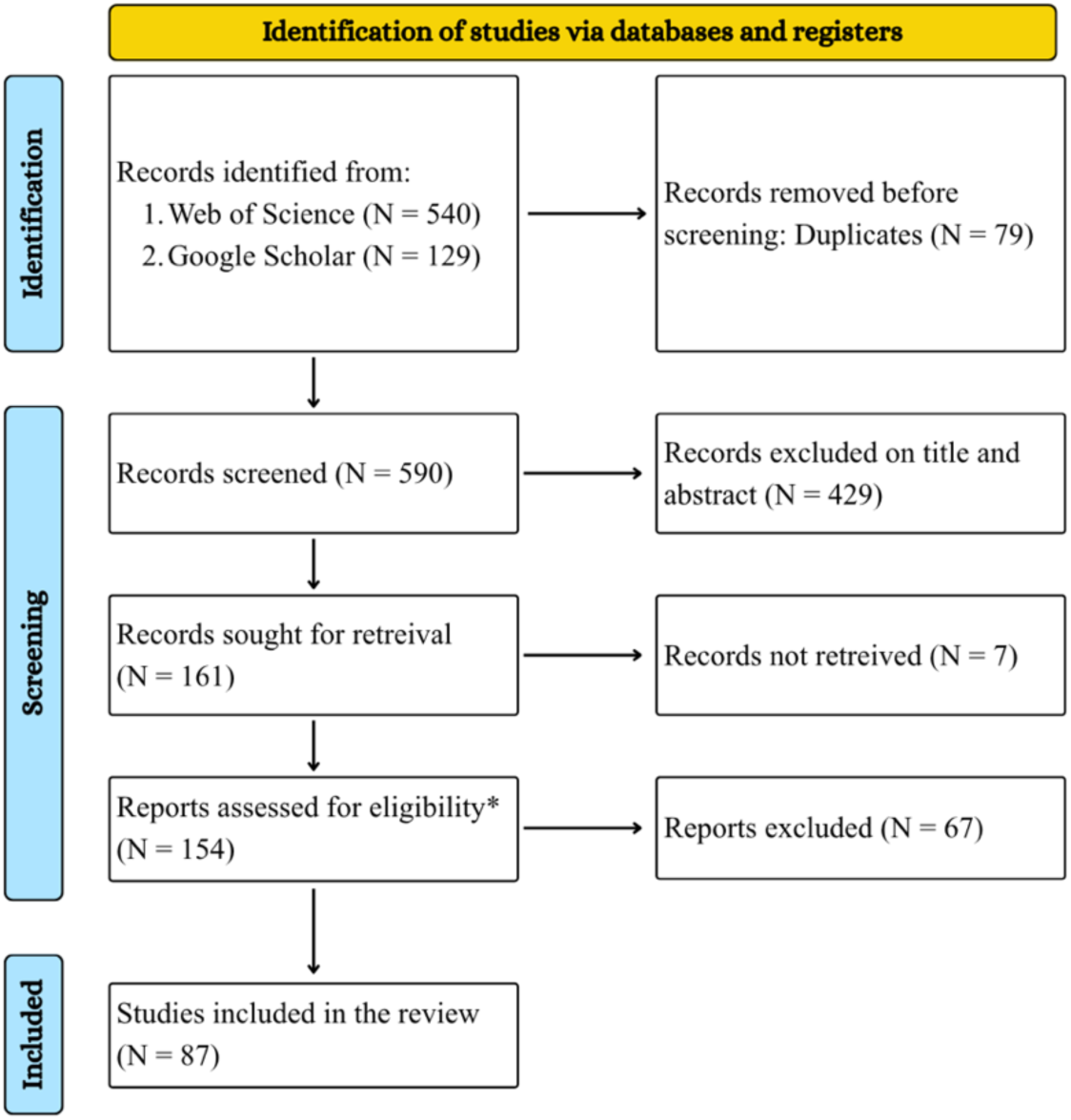
PRISMA flow diagram, illustrating the systematic review process taken when investigating global emergence time in bats. *Eligibility criteria: (1) direct report of the mean emergence time; (2) availability of raw emergence time data available from the publication, supplementary materials or on correspondence with the author.

### 2.2 Trait dataset

To investigate the impact of various roost demographics, the following variables were extracted/calculated from the emergence time sources and included in the final analyses (table 1):

1. Mean emergence time (minutes) – the first response variable, defined as the mean time since sunset for a colony to emerge from their roosts. For studies reporting raw emergence times across a field season, we averaged emergence time per day over the entire sampling period, thereby accounting for seasonal shifts in sunset time.
2. Brightness at emergence (radians) – the second response variable, measured as the altitude of sun at emergence with respect to the horizon. Using the mean emergence times reported per study, we calculated sun’s altitude at emergence per roost per study-specific sampling period. We used the following R packages to calculate and compile this variable – “suncalc” (Thieurmel & Elmarhraoui, 2017) and “lubridate” (Spinu et al., 2024).
3. Latitude and longitude (°) – geographical coordinates of each roost (fig. 3), as reported in their respective studies. Latitude was retained as the primary predictor for emergence patterns, taken in its absolute form prior to analysis.
4. Diet – a categorical fixed effect, based on the focal species; categorization was done using reports from a variety of sources:

a. Insectivores – predominantly an insect-diet but also includes species that prey on small vertebrates.
b. Phytophagous – fruits, nectar, pollen or other parts of a plant.
5. Group composition – a categorical fixed effect describing the social structure of the colony being studied:

a. Maternal colonies – colony of reproducing females containing pregnant and lactating females; these roosts also contain juveniles as the season progresses.
b. Non-maternal colonies – includes colonies with non-reproductive females, males or mixed compositions. This category also includes colonies with unknown compositions (e.g., from acoustic or visual survey studies).
6. Study format – a categorical fixed effect accounting for the variation in data collection in respective studies. Since the format of recording emergences differed between studies, it created a gradient of precision that might have impact final recorded emergence times:

a. Passive acoustic surveys – data is collected in real-time by detectors placed either right outside roosts or in the near vicinity of roosts, i.e., bats’ foraging grounds.
b. Visual count surveys – observers count and record bats exiting a known roost in real-time.
c. Radio-tagged individuals – exits and flight patterns from across the night are recorded by radio-tags attached to individuals.
7. Wing loading (N/m^2^) – a continuous variable defined as species specific averages of the ratio between body mass and wing area. This variable was included to account for variation in morphology and flight performance between species. It was extracted from a variety of published sources (see, suppl.) instead of the primary literature utilized for the emergence times.

**Fig. 3:**
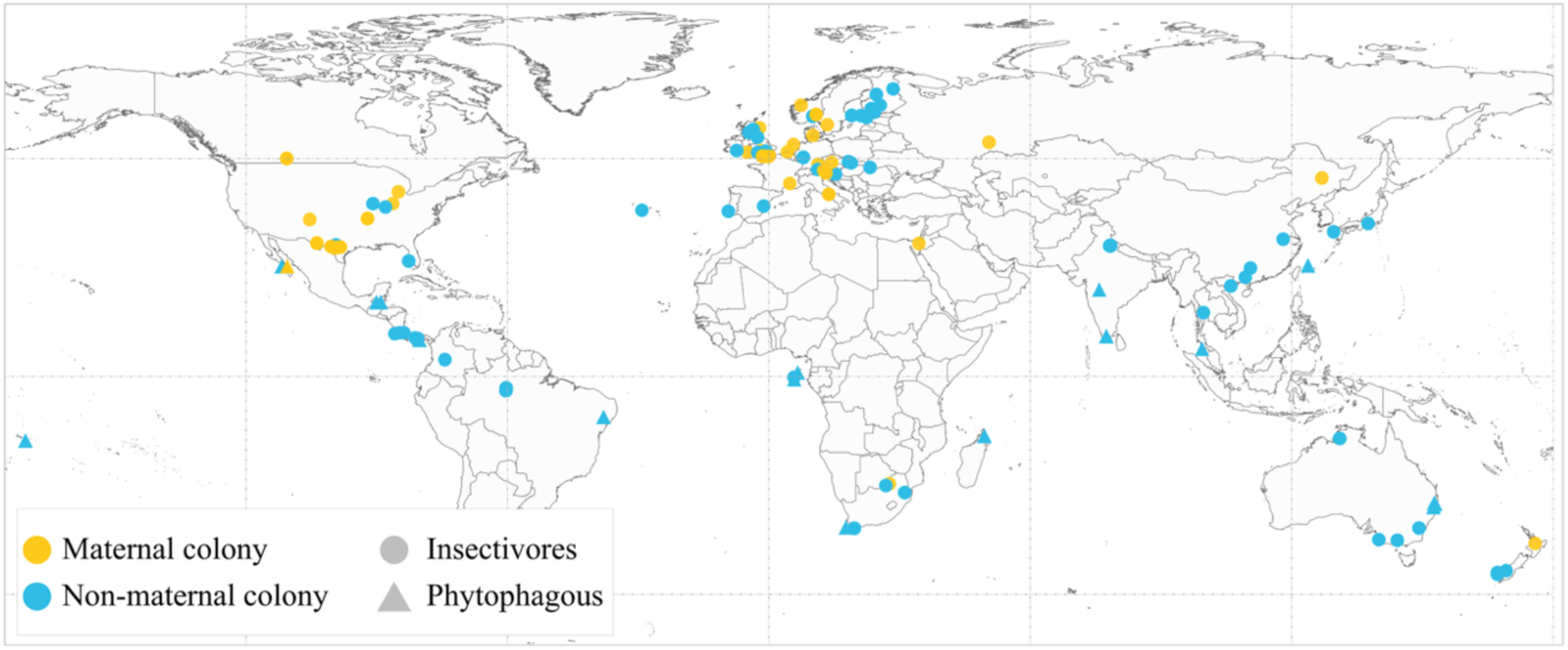
Locations of roosts included in the current analysis, as extracted from the coordinates reported in the respective sources.

Seasonality of each study was noted but later dropped due to the lack of variation it presented, i.e., nearly all studies took place during months conducive for bat activity in the respective regions.

### 2.3 Statistical analyses

All analyses for this study were conducted in R (ver 4.4.1), using Markov Chain Monte Carlo generalised linear mixed models (MCMCglmm) from the “MCMCglmm” package (Hadfield, 2024). We maintained latitude as the constant primary predictor against our responses - emergence time and brightness at emergence. Besides latitude, group composition, study format, diet and wing loading were included as additional predictors to account for roost and individual level differences. However, initial analysis revealed that wing loading did not significantly contribute to our candidate models. Based on this, we removed wing loading from our final dataset. Since wing loading was only reported for 77 species (see suppl.) dropping the variable allowed us to increase final species sample size. By dropping wing loading our dataset then contained complete cases from 100 species.

To account for shared ancestry in these models, we used a Chiropteran phylogenetic tree. This tree was extracted from the mammal phylogeny published in PHYLACINE 1.2.1 (Faurby et al., 2019) and contained 1,162 bat species. All phylogenetic data was processed and visualised using the R packages “ape” (Paradis et al., 2024) and “phytools” (Revell, 2025). Prior to including the phylogeny in the model, we cross-referenced the tree tips with our dataset, including only species that were present across both structures. Post this pruning, the final dataset used for our analyses contained 92 species from 13 families (fig. 4).

**Fig. 4:**
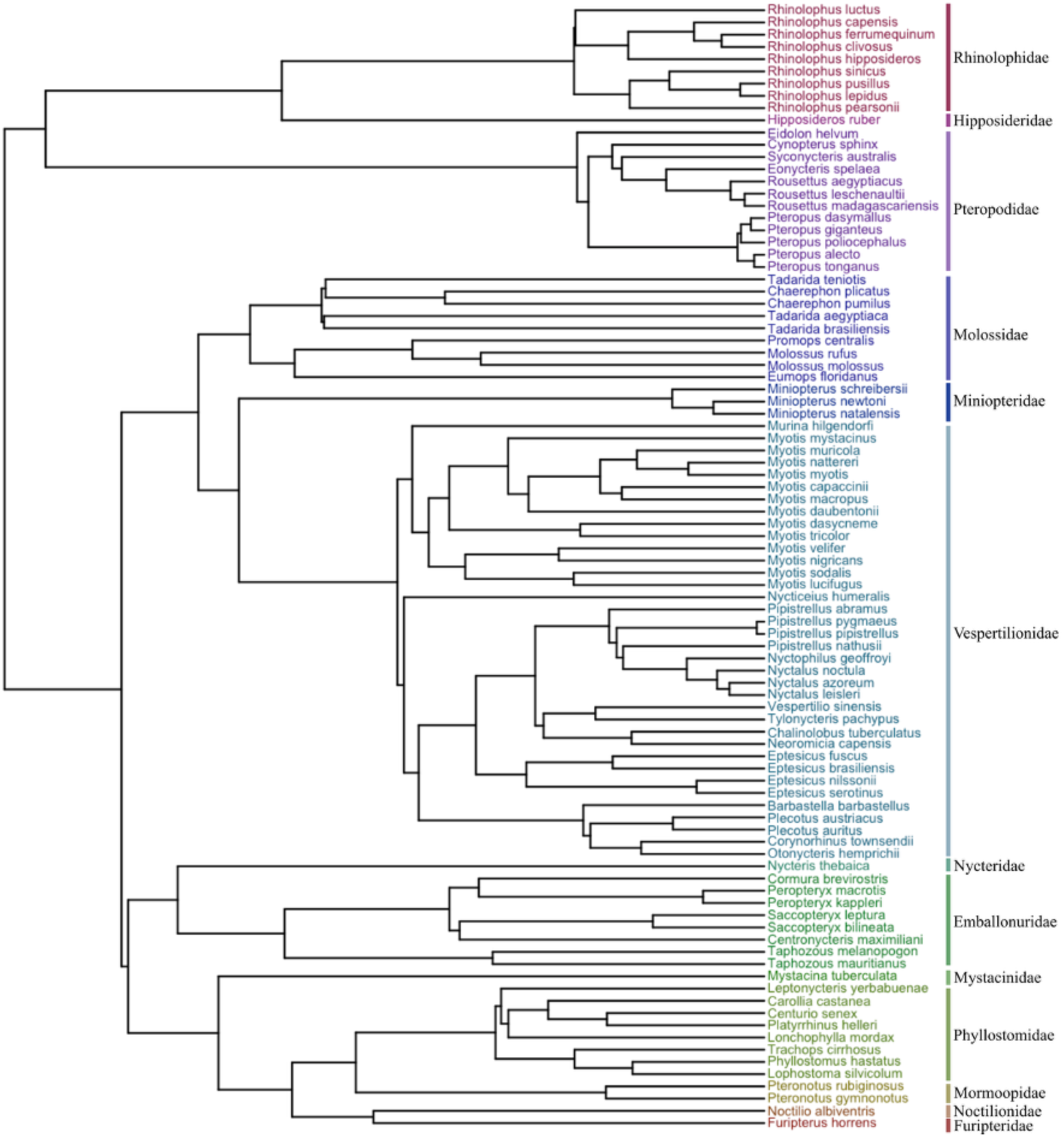
The phylogeny used to account for effect of ancestry on emergence times in bats. A total of 92 species from 13 families are included in this tree, as denoted by the tip labels and notations respectively. The length of this tree is unknown.

The MCMCglmm models were run for 130,000 iterations with a burn-in of the first 3000 iterations and a sampling rate of 50 to reduce autocorrelation. The trimmed chiropteran phylogeny was included in all models as a random effect, modelled on a weakly informative inverse-Wishart prior (variance, V = 1, degrees of beliefs, v = 0.002). The residual variance was modelled using an equivalent weak prior, allowing us to estimate posterior distributions for phylogenetic effects while accounting for uncertainty in the variance components. Each model was run on two independent chains with random starting seeds to assess convergence, with each chain resulting in an effective sample size of at least 2000. All models converged successfully using the Gelman-Rubin criterion, <1.1 (Brooks & Gelman, 1998).

Model fits were compared using DIC (Deviance Information Criterion) values as reported in the MCMCglmm models. Those with the lowest DIC scores were retained as candidate models, and those within ΔDIC ≤ 2 were considered to have comparable support. After confirming convergence, posterior predictive distributions were generated using post-burn-in samples, pooled across the chains. Final predictions were obtained by marginalising over categorical variables not of interest, allowing visualisation of the changes in the response variables against latitude.

Finally, we calculated Pagel’s lambda (λ) to assess the strength of our phylogenetic signals in the response traits. Values for Pagel’s lambda range from 0 (no phylogenetic signal) to 1 (strong phylogenetic signal consistent with Brownian motion evolution).

### 2.4 Statistical validation – accounting for variances

To account for potential daily variation that may have been lost as a consequence of using reported mean emergence times across studies, we replicated our analyses in 2 subsets of the data. The first subset comprised 15 long-term studies that provided raw data for their entire study periods. This subset spanned 22 species and a range of latitudes. Using the daily roost exit times reported in each study, we calculated the corresponding values of luminosity at emergence. This data was then fit to our larger dataset’s best-fit models. Similarly, our second subset of studies were those that reported standard error alongside emergence times. We fit this, along with the measurement error variance (mev), into our candidate models for emergence times. Since brightness at emergence was derived from emergence times, SE values were absent for it. Therefore, this subset analysis was carried out on emergence times from 59 studies, spanning 67 species.

### 2.5 Case studies: emergence patterns in predator-free zones

Despite bats being primarily nocturnal, there are several observations of bats flying during the day around the world (Chua & Aziz, 2019; Hirakawa, n.d.; Russo et al., 2007; Speakman, 1991; Vivas-Toro & Murillo-García, 2020). While the explanations for these infrequent incidents vary from individual energetic needs (Frick et al., 2012) to canopy cover (Russo et al., 2011), the lack of natural predators in certain island ecosystems has been observed to allow for consistently earlier evening emergences (Peel et al., 2017; Speakman & Irwin, 2003). Here, we aimed to quantify and compare the difference in emergence time and brightness between bat populations without natural predators versus exit patterns in related species at similar latitudes where natural predators were present.

The data for this analysis was taken from studies that explicitly state emergences, either in the raw format (exit onset and end) or as mean exit times. However, due to few predator-free environments, only two studies were considered here: one in the temperate region and one in the equatorial region. The temperate region study reported emergence times in *Nyctalus azoreum*, an insectivore bat species on the Azores-island (38.53° N, Speakman & Irwin, 2003). We compared this species against two insectivorous species: *Tadarida teniotis* from Portugal (38.3° N, Marques et al., 2004) and *Nyctalus leisleri* from Ireland (52.14° N, Shiel & Fairley, 1999). In the equatorial region we used data on the frugivorous species *Eidolon helvum* from São Tomé and Príncipe (0.05° N and 1.58° N, Peel et al., 2017) as a predator-free population (see Araújo-Fernandes et al., 2025). We compared exit data of this species against data collected from other frugivores, *Pteropus medius* and *Cynopterus sphinx* from India (9.97° N, Elangovan et al., 1999; 9.93° N, Murugavel et al., 2023) as well as *Eonycteris spelea* from Thailand (7.07° N, Acharya et al., 2015). See Fig. 6a for map of locations.

We ran a total of four two-way ANOVAs testing exit times (minutes since sunset) and brightness at emergence (rad) as response variables for each region (temperate and equatorial). Each model included predator presence and species identity as fixed predictors.

## Results

We ran 9 models per response variable, with latitude as the primary predictor in 8 of them (1 intercept model). Using the DIC scores reported in each model, we retained models with the lowest scores as our candidate/best-fit models (table 2).

**Table 2:** Model selection support, based on DIC scores, written in decreasing order of support.

| Emergence time models | ΔDIC | Brightness at emergence models | ΔDIC |
| --- | --- | --- | --- |
| **Latitude + group composition + study format** | **00.0** | **Latitude + group composition + study format** | **00.0** |
| Latitude * diet + group composition + study format | 03.0 | Latitude * diet + group composition + study format | 02.7 |
| Latitude + study format | 10.4 | Latitude + study format | 08.3 |
| Latitude * diet + study format | 13.9 | Latitude * diet + study format | 11.2 |
| Latitude + group composition | 14.4 | Latitude + group composition | 12.0 |
| Latitude * diet + group composition | 16.8 | Latitude * diet + group composition | 14.4 |
| Latitude | 27.3 | Intercept only | 18.2 |
| Latitude * diet | 29.6 | Latitude | 19.4 |
| Intercept only | 30.8 | Latitude * diet | 21.8 |

As per these results, the most prominent variables responsible for emergence patterns were latitude, group composition within roosts and study format. Diet was note seen playing a crucial role in determining emergence patterns.

### 3.1 Emergence time

Emergence timings were best explained by an additive combination of latitude, roost composition and the study format. The results revealed latitude as a significant predictor of emergence time. Additionally, differences in emergence times between maternal and non-maternal colonies as well as different study formats were also noteworthy (table 3a). The model reported a moderately weak phylogenetic signal for emergence patterns (λ = 0.40, 0.19 – 0.61).

**Table 3:** Results from the best-fit models for emergence patterns. The reference levels for the two fixed effects in the model were taken as maternal colonies and acoustic studies.

|  | Posterior mean | 95% CI | pMCMC |
| --- | --- | --- | --- |
| (a) Emergence time |  |  |  |
| Intercept | 2.17 | -19.84 – 22.80 | 0.836 |
| **Absolute latitude** | **0.79** | 0.47 – 01.14 | **<4e-04** *** |
| **Non-maternal colony** | **15.72** | 5.83 – 25.46 | **0.003** ** |
| Radio-tag studies | -3.51 | -14.51 – 07.38 | 0.530 |
| **Visual counts and survey studies** | **-22.27** | -33.32 – -12.22 | **<4e-04** *** |
| (b) Brightness at emergence |  |  |  |
| Intercept | -0.06 | -0.15 – 0.01 | 0.12 |
| Absolute latitude | -0.00 | -0.00 – 0.00 | 0.23 |
| **Non-maternal colony** | **-0.05** | -0.08 – -0.02 | **0.01** ** |
| Radio-tag studies | -0.01 | -0.05 – 0.03 | 0.75 |
| **Visual counts and survey studies** | **0.06** | 0.02 – 0.10 | **0.00** ** |

As per the model results, emergence time at the equator (0°), for a maternal colony, occurs at sunset. Per unit increase in latitude (∼1° increment) results in a ∼ 1 minute delay in emergence, i.e., roost exit is delayed at higher latitudes (β = 0.79, 95% CI = 0.47 – 1.14 mins). After accounting for the influence of phylogeny and latitude, maternal colonies showed significantly earlier emergences, exiting 15.72 minutes (95% CI = 5.83 – 25.46 min) before non-maternal colonies at the same latitude (fig. 5a). Lastly, the differences observed among study formats indicates that visual surveys reported emergences 22.27 minutes earlier than acoustic studies (see suppl. S1).

**Fig. 5:**
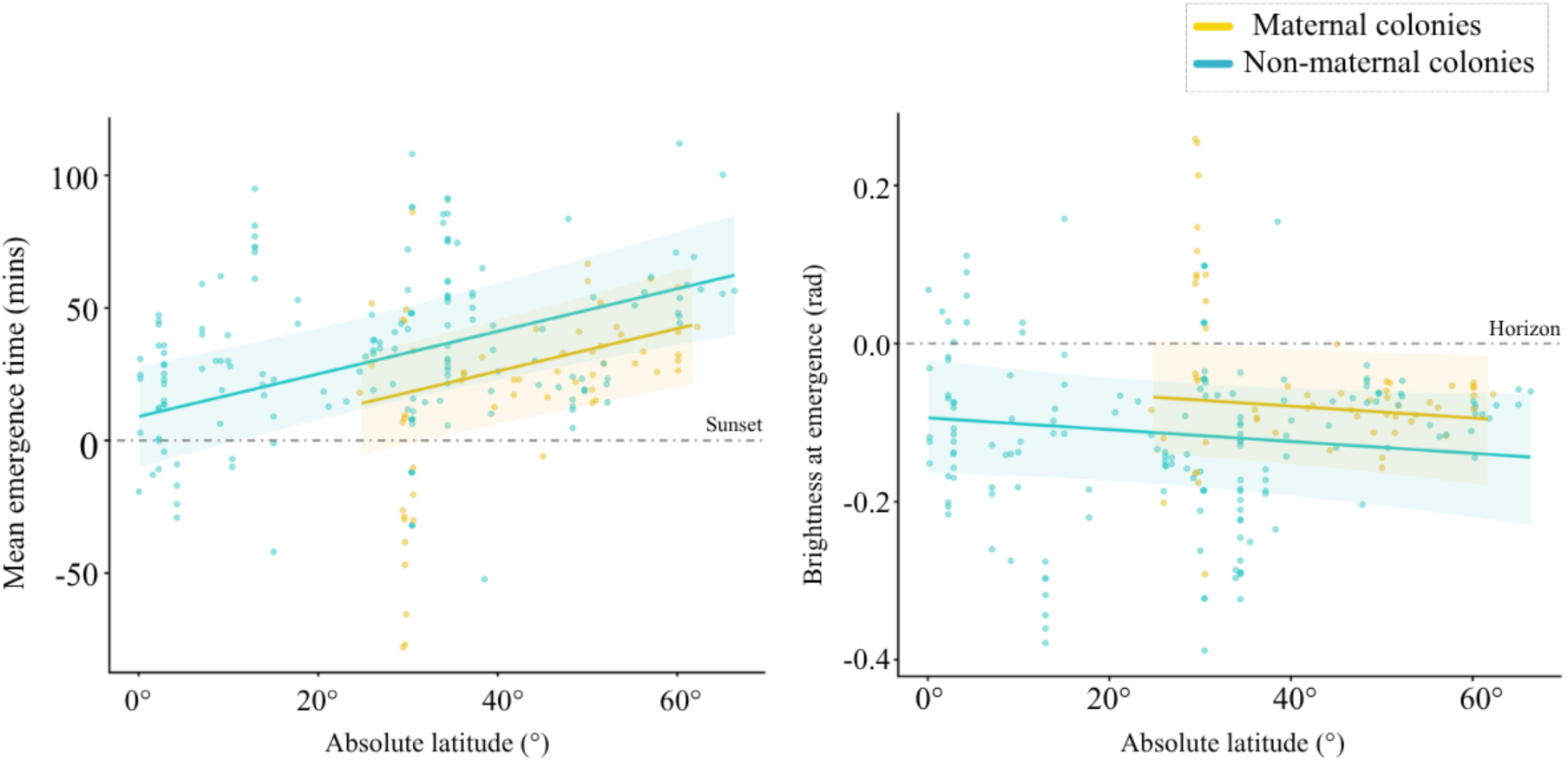
Latitudinal variation in (a) emergence time (mins) and (b) brightness at emergence (rad), as per the candidate models. Points indicate to the raw spread of data (model parameters unaccounted for); grey dot-dashed lines represent sunset and horizon respectively.

### 3.2 Brightness at emergence

Brightness at emergence, a derivate of emergence times, mirrored its results and was best explained by an additive combination of latitude, roost composition and study format. However, latitude did not play a significant role in this model, i.e., brightness at emergence was conserved across latitudes (table 3b). Lastly, this trait possessed a moderate phylogenetic signal, λ = 0.46 (0.26 – 0.65).

According to the model predictions, emergences were concentrated after sunset (fig. 5b), often occurring between sunset and nautical twilight (-0.2 rad or -12°). Maternal colonies continued to emerge under slightly brighter conditions than others (β = -0.05, 95% CI = -0.08 – -0.02 rad). Lastly, the variation in emergence reported by different study formats continued to persist (see suppl. S1).

### 3.3 Statistical validation – accounting for variances

Per our subset analysis, we found that daily data from long term studies showed similar emergence patterns as the compiled dataset. Here, higher latitudes showed prolonged periods of emergences, but the brightness at exit remained constrained to the same threshold seen in the larger model output. Similarly, accounting for the measurement variance in emergence times did not significantly alter the candidate model’s results. These analyses validate our findings, further indicating that daily variation in emergence times did not alter the overall model outputs obtained in this study. The results of these analyses can be found in the supplementary material (see suppl. S2 and S3).

### 3.4 Case studies: emergence patterns in predator-free zones

The two-way ANOVAs for the temperate zone revealed that *N. azoreum* emerged significantly earlier (F_1,11_ = 18.86, p = 0.001) and under significantly brighter conditions (F_1,11_ = 17.31, p = 0.002) compared to *N. leisleri* or *T. teniotis* (Fig. 6b – c). Similarly, *E. helvum* emerged significantly earlier (F_1,8_ = 135.67, p < 0.001) and under brighter conditions (F_1,8_ = 135.67, p < 0.001) compared to *P. medius*, *C. sphinx* or *E. spelea* (Fig. 6d – e). Both *N. azoreum* and *E. helvum* inhabit predator-free environments, and emerged consistently before sunset, while nearby or related species in systems with predators generally emerged after sunset.

**Fig. 6:**
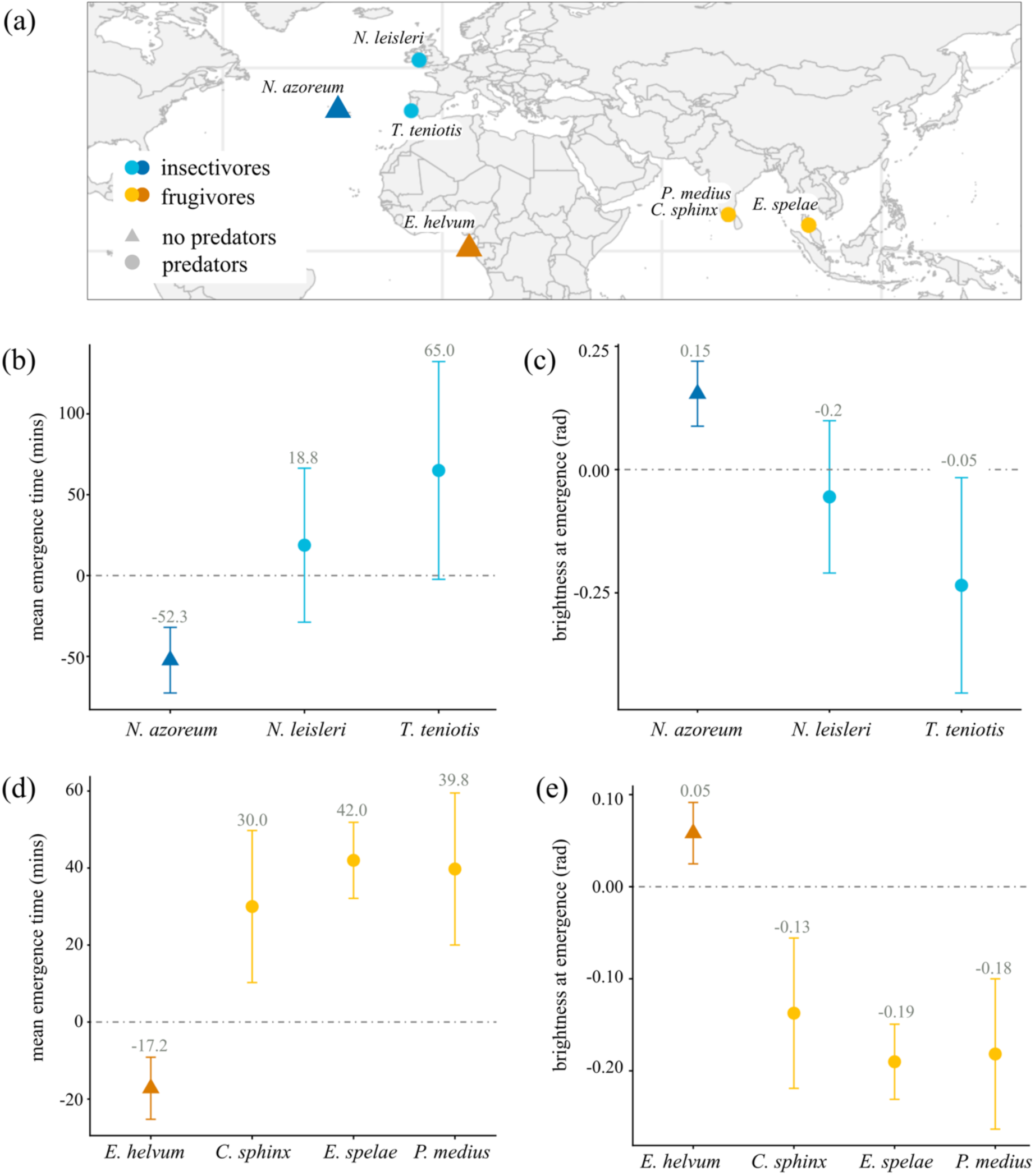
(a) Location of study sites included in this case study. A comparative analysis of exit patterns in the temperate zone showing differences in (b) mean emergence time (mins from sunset) and (c) brightness at emergence (rad) for *N. azoreum* v. related or nearby species. Similarly, a second analysis on exit patterns in the tropical zone, highlighting the variation in (d) mean emergence time (mins from sunset) and (e) brightness at emergence (rad) between *E. helvum* v. related or nearby species. The grey numbers above species error bars represent the estimated species-specific means as per respective models. The grey dot-dashed line indicates sunset / horizon.

## Discussion

Our results indicate that emergence occurs within a fixed sun altitude, with no significant variation across latitudes. Subsequently, the time of emergence (i.e., time since sunset) is delayed with increasing latitude, where the temporal delays accommodate longer twilights at higher latitudes. This contradicts our hypothesis of species inhabiting higher latitudes foraging under brighter conditions. In addition, we found a pattern of early emergences in maternal colonies, as we expected. Furthermore, though highly plastic, emergence behaviours showed a moderate effect of phylogeny. We also detected an effect of study format (i.e., acoustic surveys, visual counts, radio-tagging) where studies using visual surveys reported earlier emergences than those that used passive acoustic surveys or radio-tagged individuals. This disparity potentially arises as a result of passive acoustic recorders being placed further away from roosts, thereby recording bats with a slight temporal lag.

The axial tilt of Earth causes prolonged periods of twilight (dusk and dawn) at temperate latitudes during summer, i.e., the period of activity for bats (commonly insectivores) inhabiting these regions. Therefore, for a given time interval after sunset and before sunrise, ambient conditions of brightness are lower at tropical latitudes than at temperate ones (see, fig. 1b). Our results indicate that roost emergence across latitudes occur at a conserved light thresholds – around nautical twilight, when the sun drops ∼ 11.5° below the horizon. Because the sun sets at a shallower angle at temperate latitudes, it takes longer for the sun to reach this threshold. This is well reflected in our finding on temporal delay in exits of bats across latitudes, with mean emergence time at 60° N occurring nearly 44 minutes after sunset.

The observed pattern of roost emergence occurring around a certain level of ambient light, as evident across both latitudes and the bat phylogeny, suggests a strong selection pressure on the behaviour. For insectivorous bats, this strict pattern of roost emergence can be explained through temporally fluctuating costs and benefits, such as predation risk from diurnal raptors, mobbing behaviour by larger birds, risk of overheating, and insect availability (Speakman et al., 2000). Because predation is ultimately fatal to the prey, selection should act upon an anti-predatory trait that decreases the risk of becoming prey and therefore provides a fitness advantage (Lind & Cresswell, 2005). Indeed, risk of predation influences decision-making in various animals, particularly with respect to activity levels and active periods (Creel & Christianson, 2008; Laundre et al., 2010; Lima & O’Keefe, 2013). Bats often roost in colonies with numbers of individuals ranging from tens to thousands, in some cases even millions of individuals. Emergence from these roosts often occurs from a single exit, in close succession, thereby presenting opportunistic diurnal predators, such as birds of prey, with increased predatory success rates (Fenton et al., 1994). However, for visual-hunting raptors this success rate rapidly dwindles with decreasing light levels (Mitkus et al., 2018). Therefore, roost emergence presents a high-risk situation where delaying emergence to a threshold light intensity is beneficial for survival.

However, the absence of diurnal avian predators allows bats to exploit the crepuscular peak completely by emerging before sunset. This pattern has been observed occurring consistently on the Azores islands (Leonardo & Medeiros, 2011; Moore, 1975; Speakman & Irwin, 2003) as well as the São Tomé and Príncipe islands (Peel et al., 2017). Results from our case study highlight the pre-sunset timelines and brightness at exit of *N. azoreum* and *E. helvum*, the two focal species inhabiting the two islands respectively. The patterns of earlier and brighter exits in the two species become more evident when compared to closely related species that occupy similar latitudinal ranges. Thus, it appears that predator-prey trophic interactions, across dietary guilds, may be acting on the selection for roost departure, limiting it to a narrow range of light levels (Arndt et al., 2018; Duvergé et al., 2000; Fjelldal et al., 2025; Frafjord, 2021; Lima & O’Keefe, 2013).

The similarities in emergence patterns across dietary guilds reflect the influence of foraging opportunities on bats’ behavioural decisions (Jones & Rydell, 1994; Welbergen, 2008). For insectivorous bats, the prey display diel patterns of activity, with a peak in activity during the crepuscular period (Gårdman et al., 2026). Similarly, for phytophagous bats, the availability of fruit and nectar declines as the night progresses (Bumrungsri et al., 2013; Gould, 1978; Sritongchuay et al., 2008). Therefore, to maximise foraging success, bats across dietary niches should attempt to time their evening emergence to overlap with this period of maximum food availability (Acharya et al., 2015; Cano & Murillo-García, 2021; Lilley, Fjelldal, et al., 2026). This was mirrored in our results, where diet was not significant in explaining global emergence patterns.

Furthermore, our results indicate the threshold of brightness for emergence is not only observed across dietary guilds, but across the order Chiroptera as a whole. Previous studies have attributed the variation in emergence strategies to ecological conditions (Geipel et al., 2019; Russo et al., 2007), predatory pressures (Arndt et al., 2018; Lima & O’Keefe, 2013) and reproductive states (Fjelldal et al., 2025; Lilley, Fjelldal, et al., 2026). Building on these, our findings also report moderate phylogenetic signals associated with these emergence behaviours, suggesting that ecological factors interact with species’ evolutionary history. A potential explanation for the effect of phylogeny could be found in physiology related to the bat wing. The combination of wing morphology and body mass in bats influences whether they are agile and fast fliers, or more manoeuvrable, but slow flying (Norberg & Rayner, 1987). Flight speed may impact the chances of being caught by raptorial birds and has therefore been associated with species-specific emergence times in bats, with slow flying species emerging later than more agile species (Jones & Rydell, 1994). Wing loading can be used as a measure of flight speed, which we considered in our analyses: however, we did not detect any effect of wing loading on emergence patterns in early models. Since the wing loading values used here were species-specific, primarily describing non-reproductive adult males, it is likely to have diminished its effect as an allometric control of morphology and flight performance.

Lastly, our statistical validation results reflected the results from our primary analyses. Species altered their emergence times to accommodate the shift in brightness over the entire study period, a pattern present irrespective of latitudes. These results are consistent with other species-specific studies that have reported populations tracking changes in sunset times across seasons at different latitudes, altering their emergence times to fit the changing conditions of light (Fjelldal et al., 2025; Frafjord, 2021; Welbergen, 2008). Overall, our results regarding threshold brightness for emergence likely reflect general patterns, even if fine-scale variances are not captured.

Based on these findings, we note bats’ emergence behaviours to be a function of latitude. Furthermore, it appears to be a trait that is moderately conserved across the phylogeny. This suggests that timing of emergence has certain fitness benefits, indicating that this behaviour could be under selection. Although representing a small fraction of the taxa, our findings advance current knowledge regarding factors that shape emergence strategies in bats. Future studies that measure light levels at emergence will be critical in substantiating these patterns. These results highlight the importance of considering latitudinal gradients in photoperiod and light intensities when investigating animal behaviours that are governed by circadian or circannual rhythms.

## Acknowledgements

We would like to thank Will Morris for his input on the modelling approach. SS acknowledges Erasmus+ for initial salary and mobility funding. RC acknowledges salary support from the Deutscher Akademischer Austauschdienst (DAAD) (at the time of data collection) and by the Centre for Wildlife Studies (CWS), India, and fieldwork funding from the Rufford Foundation and Wildlife Acoustics.

