## Supplementary material for "The geography of nocturnality – emergence patterns of bats on a latitudinal gradient": S1 S2 S3

**S1 Study format variation as per model results**

The best-fit models for emergence time and brightness showed significances in the estimates generated for different study formats. As per the model results, visual counts and surveys reported emergences 24.21 minutes before passive acoustic recorders. This disparity highlights the differences in grain of precision in data recording formats for studies looking at emergence times.

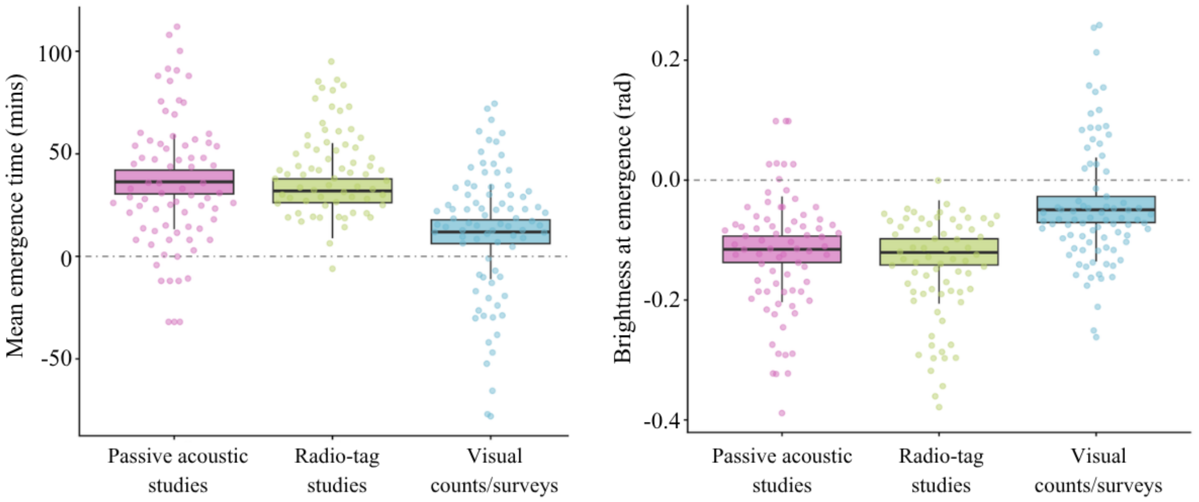


Fig. S1: Best-fit model results of (a) emergence time and (b) brightness as recorded by different study formats included in this study. The points represent the spread of the raw dataset. The grey dotted line represents sunset and horizon respectively (0 mins to / from sunset and 0.0 rad).

**S2 Statistical validation using long term data**

The subset of data used for this analysis comprised of 4 maternal colonies and 13 non-maternal colonies, and only acoustic and radio-tag study formats, i.e., no visual surveys. The model results mirrored our reported model results, with colonies at higher latitudes delaying their emergence times (table S2a). There was a stronger phylogenetic signal observed for this model; however, this is likely an artifact of a small species sample pool, resulting in seemingly stronger signals.

Similarly, the results from the brightness at exit model showed too mirrored the candidate model from the larger dataset. Species across latitudes constrained their emergences between threshold limits of brightness on a daily scale (table S2b). However, since the species pool size was small, we were unable to run with it with a phylogeny.

Table S2: Results for emergence patterns on long term studies across latitudes

| | **Posterior mean** | **95% CI** | **pMCMC** | |
|---|---|---|---|---|
| (a) Emergence time | | | | |
| Intercept | 17.18 | 0.46 – 37.29 | 0.043 | * |
| Absolute latitude | **0.14** | 0.06 – 00.20 | **<0.007** | ** |
| Non-maternal colony | **9.41** | 6.95 – 11.83 | **<0.007** | ** |
| Radio-tag studies | -24.24 | -49.98 – -02.64 | 0.057 | • |
| (b) Brightness at emergence | | | | |
| Intercept | **-0.01** | -0.02 – -0.01 | **<0.007** | *** |
| Absolute latitude | **-0.00** | -0.00 – -0.00 | **<0.007** | *** |
| Non-maternal colony | **-0.06** | -0.07 – -0.06 | **<0.007** | *** |
| Radio-tag studies | **-0.08** | -0.08 – -0.07 | **<0.007** | *** |

Since the data here largely relies on passive acoustic recorders set at a distance from any known roosts, instances of a bat being recorded nearly 2 hours after sunset are common. These recordings are likely foraging and other nightly activity of individuals in the area and do not necessarily denote emergence timestamps.

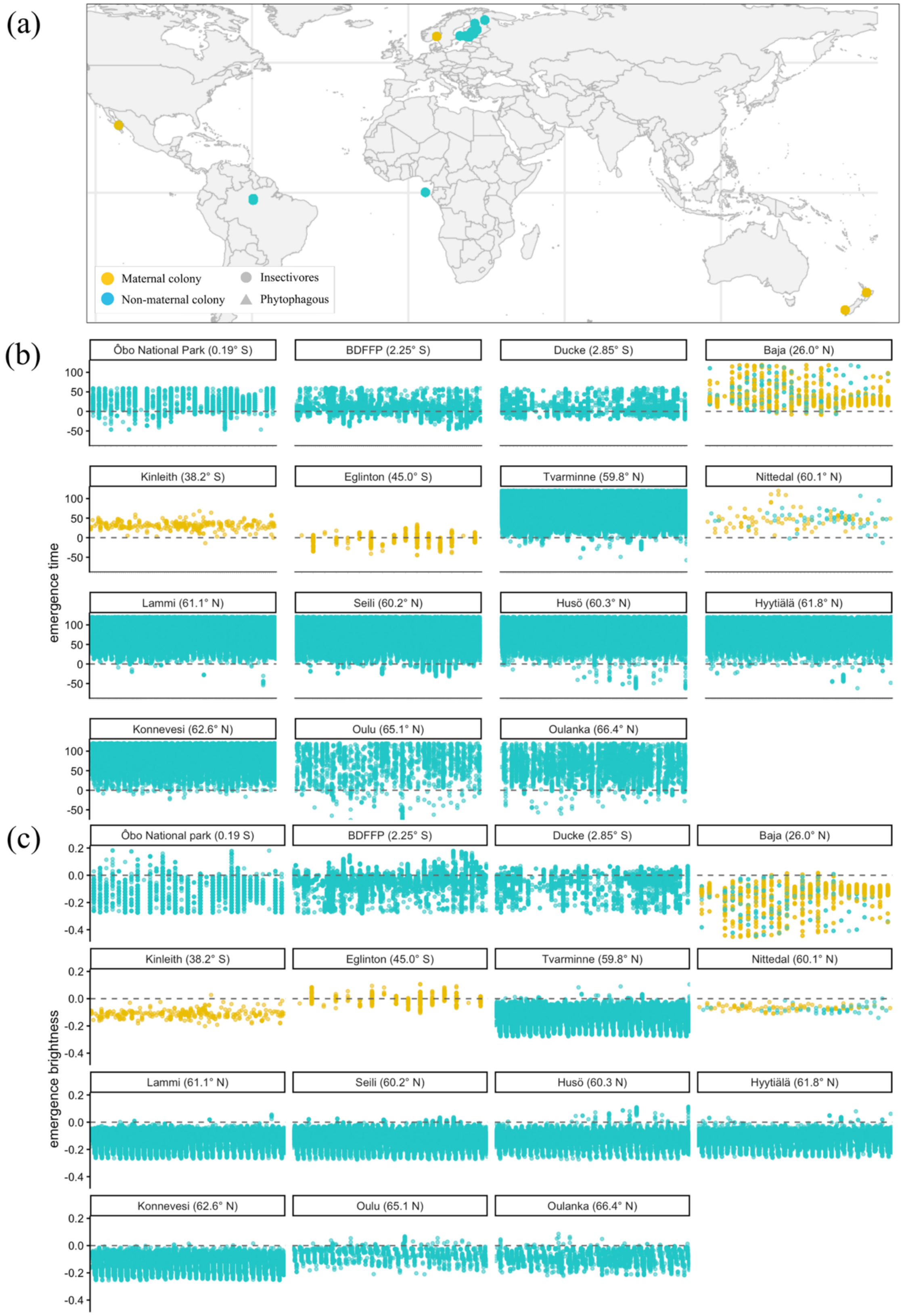


Fig. S2: (a) Locations of studies included in the long-term data analysis; (b) Daily emergence times of multiple roosts/individuals for 15 studies, from across different latitudes; (c) Daily brightness at emergence for 15 long term studies from different latitudes. Here, each data point represents one radio-tagged individual or an acoustic recording. The grey dotted line represents sunset and horizon respectively.

### S3 Statistical validation using measurement error variance

The results from the statistical validation confirmed that while accounting for SE reduced the margins of 95% CIs, they did not alter the outcome, confirming that our analysis and subsequent results are not heavily impacted or diminished by unaccounted variance measures. To ensure subset size did not influence these model results, we ran this data on two models: one with measurement error variance as part of the model (table S3a) and the second without (table S3b). The two models showed similar results (fig S3), demonstrating that the lack of effect seen between the models with and without variance is not a result of the trimmed dataset.

Table S3: Results from the statistical validation with and without measurement error variance (mev).

| | Posterior mean | 95% CI | pMCMC | |
|---|---|---|---|---|
| (a) model + mev | | | | |
| Intercept | 9.04 | -8.69 – 27.36 | 0.32 | |
| Absolute latitude | **0.66** | 0.37 – 00.98 | **<4e-04** | *** |
| Non-maternal colony | **10.79** | 1.34 – 20.49 | **0.03** | * |
| Radio-tag studies | -7.18 | -18.71 – 04.88 | 0.24 | |
| Visual survey studies | **-22.64** | -34.79 – -10.54 | **0.00** | *** |
| (b) model - mev | | | | |
| Intercept | 12.43 | -8.53 – 33.63 | 0.25 | |
| Absolute latitude | **0.71** | 0.39 – 01.06 | **<4e-04** | *** |
| Non-maternal colony | **10.51** | 0.27 – 20.51 | **0.05** | * |
| Radio-tag studies | **-12.25** | -24.52 – -00.94 | **0.04** | * |
| Visual survey studies | **-28.83** | -28.83 – 15.92 | **<4e-04** | *** |

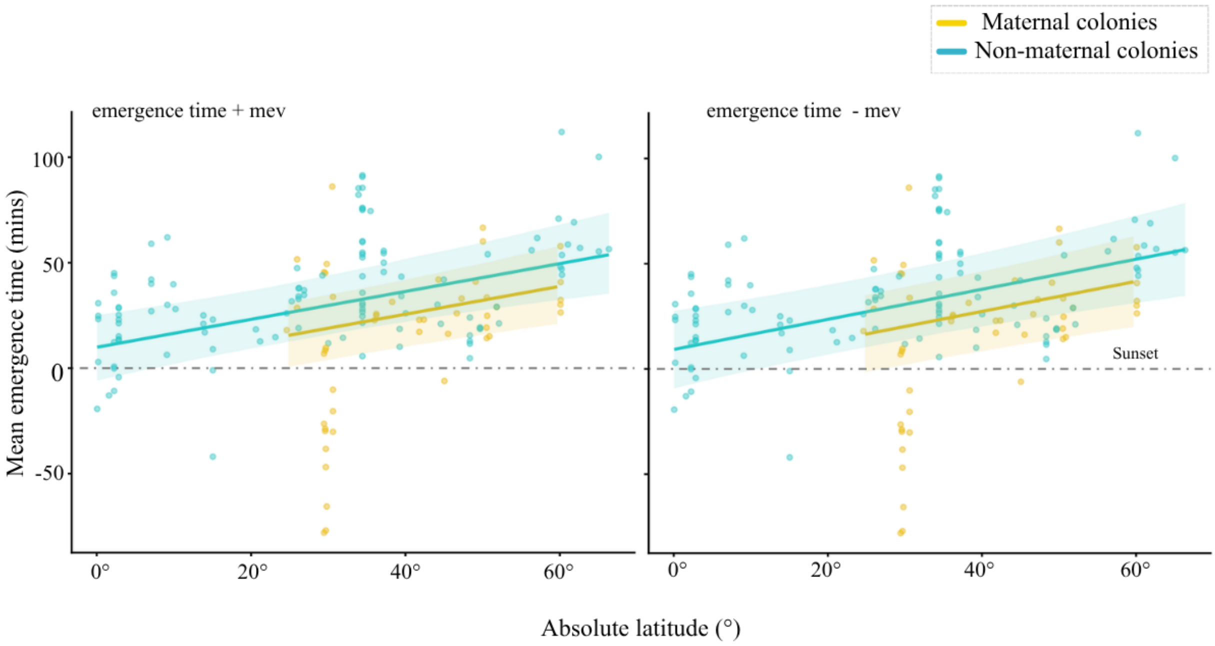


Fig. S3: Mean emergence time across latitudes, (a) accounting for measurement error variance and (b) without measurement error variance in the candidate model. The grey dotted line represents sunset / horizon. The overlayed points represent the raw emergence time data. The negative values represent emergence activity before sunset and vice-versa.
